# A velocity plan with internal feedback control best explains modulation of saccade kinematics during eye-hand coordination

**DOI:** 10.1101/2020.10.08.330647

**Authors:** Varsha V, Atul Gopal, Sumitash Jana, Radhakant Padhi, Aditya Murthy

**Author notes:** **Corresponding author**: Dr. Aditya Murthy, Centre for Neuroscience, Indian Institute of Science, Bangalore, Karnataka, India – 560012.

## Abstract

Fast movements like saccadic eye movements that occur in the absence of sensory feedback are often thought to be under internal feedback control. In this framework, a desired input in the form of desired displacement signal is widely believed to be encoded in a spatial map of the superior colliculus (SC). This is then converted into a dynamic velocity signal that drives the oculomotor neurons. However, recent evidence has shown the presence of a dynamic signal within SC neurons, which correlates with saccade velocity. Hence, we used models based on optimal control theory to test whether saccadic execution could be achieved by a velocity based internal feedback controller. We compared the ability of a trajectory control model based on velocity to that of an endpoint control model based on final displacement to capture saccade behavior of modulation of peak saccade velocity by the hand movement, independent of the saccade amplitude. The trajectory control model tracking the desired velocity in optimal feedback control framework predicted this saccade velocity modulation better than an endpoint control model. These results suggest that the saccadic system has the flexibility to incorporate a velocity plan based internal feedback control that is imposed by task context.

**NEW & NOTEWORTHY:** We show that the saccade generation system may use an explicit velocity tracking controller when demand arises. Modulation of peak saccade velocity due to modulation of the velocity of the accompanying hand movement was better captured using a velocity tracking stochastic optimal control model compared to an endpoint model of saccade control. This is the first evidence of trajectory planning and control for the saccadic system based on optimal control theory.

## INTRODUCTION

We make hundreds of saccades during our waking hours, which allows the fovea to focus on specific spatial locations. Multiple cortical structures are involved in the planning of saccadic goal, with the midbrain structure of superior colliculus (SC) being the final stage where the spatial goal is represented (Moschovichis 1987; Munoz and Wurtz 1995; Schall 2013; Sparks and Mays 2003). The brainstem saccade circuitry is thought to convert this spatial representation of the desired goal of the saccade in SC into a temporal motor command which causes the activation of oculomotor muscles and leads to the fast refocusing of the target on the fovea. This anatomical pathway has motivated theoretical studies which suggest that saccade execution is controlled based on its amplitude that is encoded in the activity of SC neurons (Jürgens et al. 1981; Robinson 1986; Scudder 1988). Neurophysiological studies on primates have shown that SC neurons increase their firing before saccades and that the amplitude and direction of the saccade are topographically encoded in the SC spatial map (Sparks et al. 1976; Wurtz and Goldberg 1972). This was further confirmed by electrical stimulation of the SC which produced fixed vector saccades (Robinson 1972; Schiller and Stryker 1972). Thus, it is well established that SC encodes a plan in the form of the desired endpoint (amplitude) of the saccade. Given this context, the dominant view is that the saccade endpoint is controlled while saccade trajectory particularly it’s velocity is just a consequence of the dynamics of the underlying neuro-muscular circuitry (Chen-Harris et al. 2008; Harris and Wolpert 2006; Sparks 2002).

Contrary to this well-established framework, a recent study showed that the SC neurons may also encode information about saccade kinematics, in addition to the amplitude and direction of the saccade. For example, instantaneous activity in SC is well correlated with instantaneous eye velocity throughout the saccade duration (Smalianchuk et al. 2018). This observation is congruent with previous work which showed the average firing rates in collicular cells are correlated with saccadic velocity (Waitzman et. al. 1991; Reppert et al. 2018; Sparks and Mays 2003; Berthoz et al. 1986). Further, reversible inactivation of SC not only affects the amplitude and direction of the saccades but also impairs the velocity of the saccade (Quaia et al. 1998). Finally, Goossens and van Opstal (2012) showed that a tight non-linear coupling between saccade amplitudes and peak velocities could be accounted for by the SC neurons (Goossens and van Opstal 2012). Hence, the traditional view that saccade generation is solely controlled by its endpoint may not be entirely justified.

In addition to these new neurophysiological findings, a series of behavioral studies showed that the peak velocity of saccadic eye movements may be modulated when happening in conjunction with reach movements. For example, saccade velocities are modulated when accompanied by hand movements (Snyder et al. 2002) or when the hand kinetics are modified by loading weights (Van Donkelaar et al. 2004) or due to reward-oriented behavior (Takikawa et al. 2002). A recent study showed that modulating the hand velocity using instructions caused modulation of the velocities of the accompanying saccades, independent of the saccade amplitude (Gopal et al. 2017). The traditional endpoint-based approach to saccade control may not be able to account for this modulation of saccade velocities that is independent of its amplitude/endpoint. Taken together, these behavioral and neurophysiological studies motivated us to revisit the traditional endpoint saccadic control approach and explore other strategies of saccade control such as trajectory tracking.

A popular approach used to understand, and model saccadic control is optimal control theory, which has successfully predicted the stereotypical behavior of the saccadic system, such as the main sequence relationship between saccade amplitude, peak velocity, and duration. Optimal control theory postulates that movements are a result of minimizing a composite movement cost. The earlier models in the optimal control framework were feedforward models which suggested that the control happens in a pre-programmed manner. Such models have suggested that the objective of the system is to achieve speed-accuracy trade-off (Harris and Wolpert 2006) or to minimize motor noise (van Beers 2008). Although nominal saccade behavior has been captured by feedforward optimal control models, they failed to explain perturbed saccade trajectories. Recently, an optimal feedback control model, which indirectly controls the displacement trajectory by predicting the instantaneous displacement with the help of a forward model has successfully explained perturbed and adaptive saccade behavior, where the desired plan at the input is still the final displacement i.e. the saccade endpoint (Chen-Harris et al. 2008; Xu-Wilson et al. 2011). Nonetheless, all these models assume endpoint control without requiring an explicit trajectory plan.

To address this caveat in the theoretical framework, we recently proposed a trajectory tracking controller for saccade generation based on desired velocity input using an optimal feedback control framework (Varsha et al. 2020, submitted manuscript). This is the first-ever optimal control model that uses an explicit plan of the desired velocity to control the saccade generation. In this study, we compared the performance of the new velocity-based trajectory tracking optimal control model of saccade in predicting saccade behavior to that of the traditional endpoint model of saccade control. We showed that the velocity-based trajectory model of the saccadic system better accounts for the saccade behavior observed during an eye-hand coordination task with explicit hand velocity modulations, but no instruction for saccades.

## METHODS

The behavioral data used in this study were taken from coordinated eye-hand experiments reported previously in (Gopal et al. 2017). The specific details of this velocity eye-hand coordination task, relevant to the current study, are discussed in brief below.

### Experimental Details

#### Human participant details

Sixteen participants (8 males and 8 females, all right-handed, corrected-to-normal vision) between the ages of 21-29 years participated in this study. The protocols of the experiment were approved by the Institutional Human Ethics Committee of the Indian Institute of Science. Written informed consents were obtained from all the participants and they were monetarily compensated for performing the task. Experimental setup details used for behavioral recording are described in (Gopal et al. 2017).

#### Task design

Briefly, the task required the participants to perform a pointing movement to a peripheral target in 3 conditions: fast velocity hand movement, slow velocity hand movement, and normal velocity hand movement. Critically, no instruction was provided for the eye, but participants almost always made an accompanying saccade with the hand movement to the target. The slow and fast trials were randomly interleaved in a block while the normal velocity trials were recorded in a separate block. Each condition was cued using different colored boxes presented at the center of the screen and written instruction at the start of each trial. A yellow cue was used for the slow trials, blue for the fast trials, and white for the normal trials. For the first 6 participants, the normal velocity block was recorded after the slow-fast block in a separate session. For the next 10 participants, normal velocity block was recorded before the slow-fast block during the same session. There was also a difference in the written cue used for the two groups. The first group had the written instruction of ‘Eye-Hand Slow’ and ‘Eye-Hand Fast’ displayed just above the cue in the slow-fast block. The second group had written instruction changed to ‘Slow Hand’ and ‘Fast Hand’ with added instruction of ‘Normal Hand’ for the normal velocity block. Since there was no systematic performance difference between the two groups of participants, the data were combined for all analyses.

The trial began with a colored cue displayed for 1500ms at the center of the screen, followed by the appearance of a white fixation box. The participant had to fixate both eye and hand for 300 ± 15 ms, following which a green target was presented either at 13⁰ or 6⁰ to the right or left of the fixation. The participant was instructed to make a pointing hand movement to the target as soon as it appeared with a velocity as specified by the color of the cue. If the participant failed to fixate for the stipulated amount of time, the trial was aborted. On successful trials in which participants reached the target with appropriate hand velocities, a tone feedback was given. Participants were given practice for ~60 trials to map their movement times for various blocks. We observed no systematic differences that are relevant to the context of this study across the 4 targets (2 eccentricities in 2 directions). Hence, we analyzed only 13⁰ horizontal rightward saccades, that accompanied the hand movements.

### Data analysis

#### Detection of saccade and pre-processing

The saccadic eye trace data was smoothened by passing through a convolution filter which averaged each data point with its two immediate neighboring points. The start of the saccade was marked when the velocity crossed 30 degrees/s, and the end of the saccade was marked when the velocity fell below this threshold. Saccades perturbed with blinks are omitted by using a velocity threshold of 700 degrees/s. Here, we analyzed only the first saccade in each trial.

#### Normalizing saccade duration

Since saccades often had different durations, we time normalized the data. The time normalization was carried out by fitting the angular displacement of the saccade using a polynomial function, which allowed the redetermination of the angular displacement at new time points. The new time points were selected by equally dividing the saccade duration in each trial into a fixed number of bins. This allowed calculating the mean across all trials as they now had an equal number of data points. The mean angular displacement and mean angular velocity at normalized time bins from 16 participants were used for all further analysis. Saccades were labeled as fast or slow or normal depending on the velocity of the accompanying hand movement.

#### Amplitude matching

The amplitude of a saccade directly affects its peak velocity. To obtain a subset of trials in which the change in peak saccade velocity between the slow and fast trials resulted from the modulation in hand velocity alone and not due to the difference in saccade amplitudes, we matched the saccade amplitudes. To do this, trials with saccade amplitudes farthest from the mean of its distribution were removed from both fast and slow trials till the distributions were not distinct from each other (Kolmogorov Smirnov test, p>0.05).

#### Statistical tests

All errors are reported in percentage as x (mean) ± y (standard deviation) %. All data were checked for normality using the Lilliefors test. If the data were normally distributed, then a two-tailed paired *t*-test was used, otherwise, a paired Wilcoxon signed-rank test was used for statistical comparisons. The level of significance was fixed at α = 0.05. Statistical significance is marked on the figures according to the following norm: *p* < 0.05 & *p* > 0.01: *, *p* < 0.01 & *p* > 0.001: **, *p* < 0.001: ***.

### Velocity-based trajectory tracking model

The goal of the model is to track the desired input in the form of saccade velocity, which is modeled as the mean velocity of all saccades to the target. A schematic of the model is given in Figure 2. The desired input is converted by the optimal controller into an optimal control signal 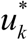, such that it minimizes the cost function given in Eq. [3]. This control signal is assumed to be corrupted by noise *ε_k_*, which is additive and signal-dependent (Harris and Wolpert 1998). The noisy control signal acts on the oculomotor plant to produce saccades. The dynamics of the plant that generates saccades given a noisy input can be modeled with the equation,

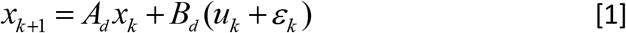

where *ε_k_=cu_k_w_k_* is the signal-dependent noise that scales with the control signal *u_k_*. The variable *w_k_* simulates the noise and is modeled as a random variable drawn from zero-mean Gaussian distribution with variance equal to one and *c* represents the noise scaling factor. The states of the system 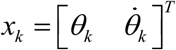 include the angular displacement *θ_k_* and angular velocity 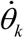 where *k* represents the time bins. The system matrices *A_d_* and *B_d_* in the Eq. [1] are obtained by discretization of the matrices *A* and *B* given by

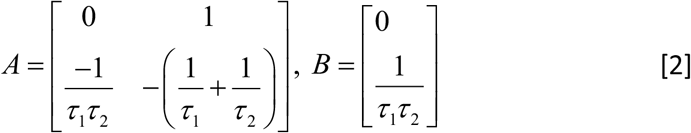

**Figure 1:**
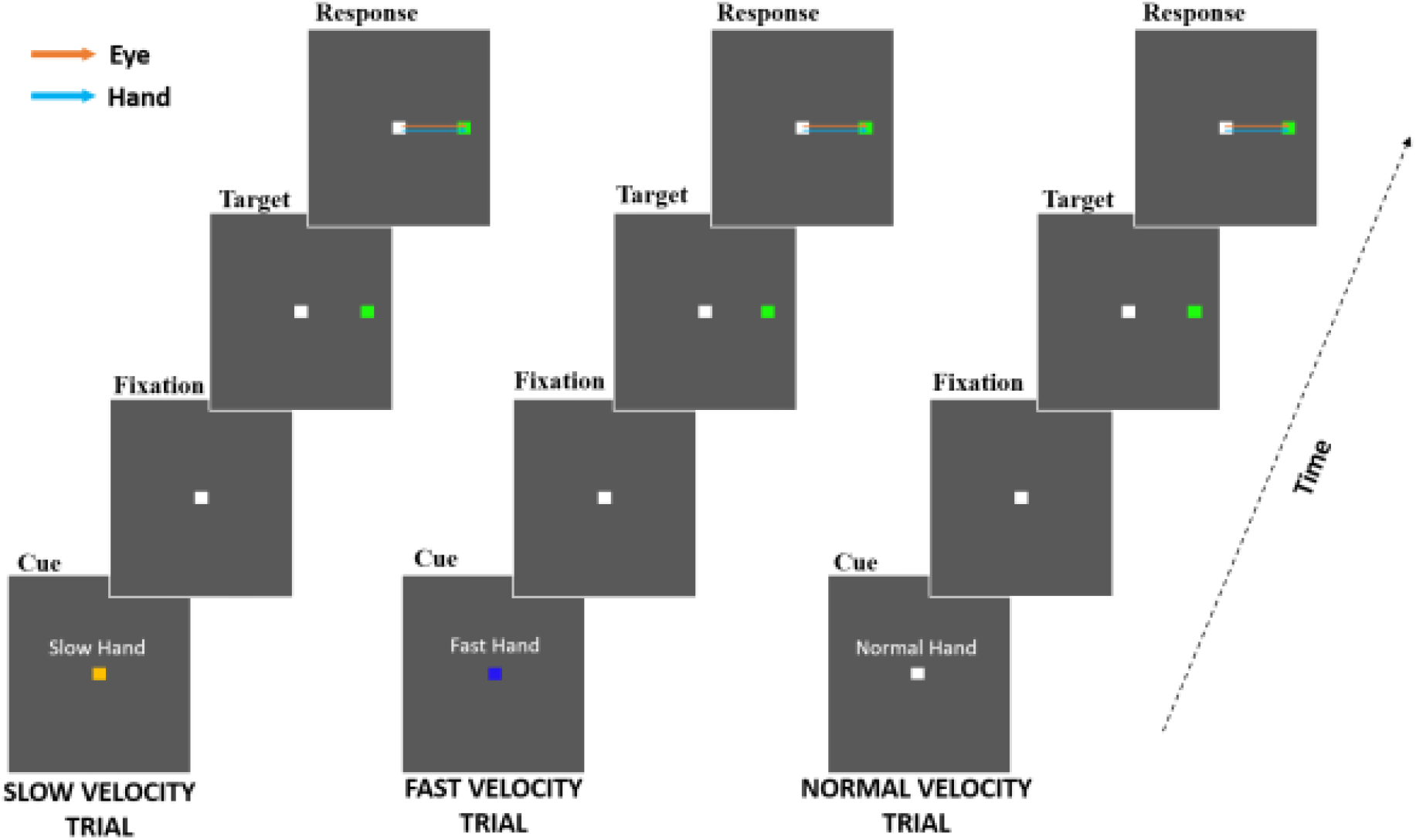
Velocity task paradigm. The colour of the square fixation box cued the participants about the desired velocity of the upcoming hand movement; yellow indicated a slow hand movement, blue indicated fast hand movement, and white indicated normal hand movement. When the peripheral green target appeared, the subject had to make pointing movement (blue trace) to the target with the appropriate velocity. No instruction was given for eye movements, but usually participants also made a saccade to the target (orange trace) (adapted from Gopal et al 2017).

**Figure 2:**
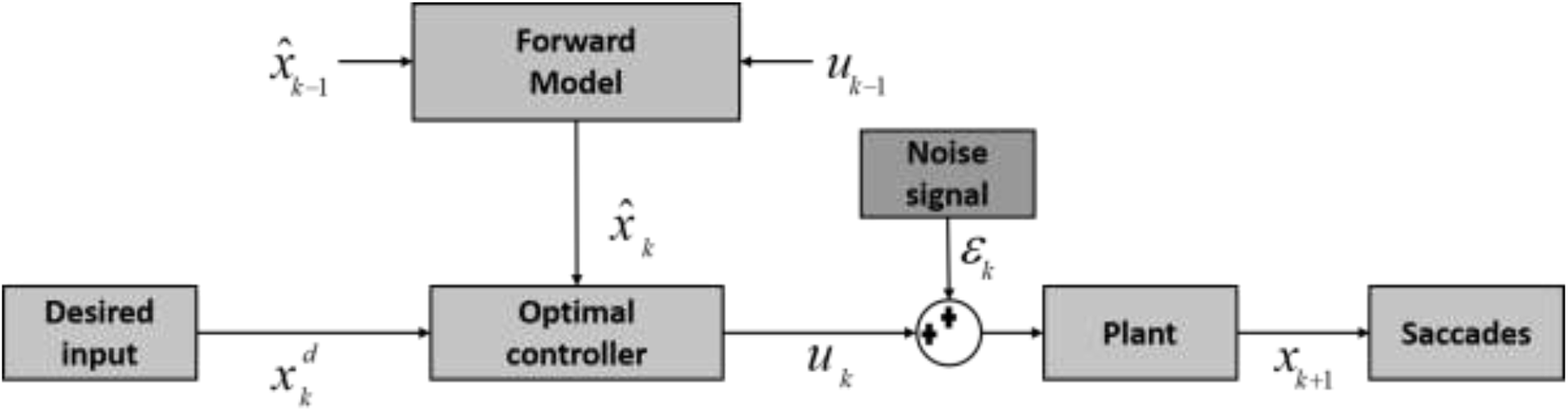
Schematic block diagram of velocity-based trajectory optimal feedback control model. The state feedback controller outputs the control signal *u_k_*. This control signal is corrupted by a noise *ε_k_*, the magnitude of which is dependent on the control signal itself. This noisy motor command causes the plant to change its state from *x_k_* to *x_k+1_*. Previous value of the control signal is used by the forward model to generate an expected value of the states.

In Eq. [2], the time constants of the plant *τ*_1_ and *τ*_2_ are assumed to be 223ms and 14 ms respectively (Robinson 1964; Harris and Wolpert 1998).

Saccadic system is not known to receive any direct external feedback signal to control its execution. Hence it is assumed that a forward model predicts the state on a moment-to-moment basis and provides internal feedback to the saccadic system (Chen-Harris et al. 2008). This forward model provides the estimate of the state 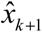 at the next instant of time based on a copy of the uncorrupted control signal and the estimate of the present state 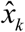. The forward model dynamics is given by 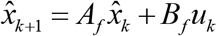 where 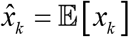. It is assumed that the forward model has evolved to perfectly imitate the oculomotor plant (i.e. Eq. [1] without the noise component) and hence for normal saccade behavior its state matrices are taken to be the same as system matrices *A_d_* and *B_d_*.

#### Optimal control signal

The most important component of the model is the optimal controller which receives the desired input and converts it into a sequence of motor commands for the oculomotor plant by minimizing the following a cost function at every step,

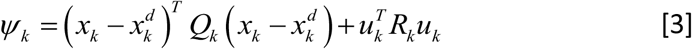

The cost function is a combination of two terms: (i) the error between the actual system state 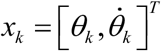 and the desired state 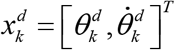 and (ii) the control effort represented by 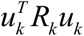. In this problem, the error weightage matrix *Q_k_* is assumed to be [0, 0; 0, *q*] with the free parameter *q*, since only the desired velocity 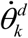 needs to be tracked. The control weighting parameter *R_k_* is fixed to one.

This problem can be solved using dynamic programming by solving for optimal value function given below over the entire movement,

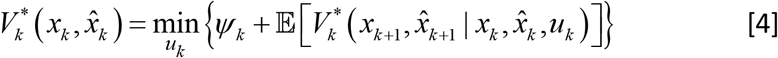

where 𝔼 represents expectation. To solve this problem, we use a value approximation approach, where the value under optimal control policy is assumed to be of the form,

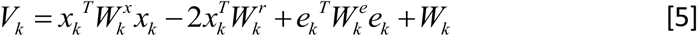

The variable 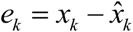 is the estimation error between the current state of the system and the estimated state predicted by the forward model. The optimal control 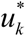 is obtained by solving Eq. [5] using value-form given by Eq. [4]. The optimal control policy is obtained as,

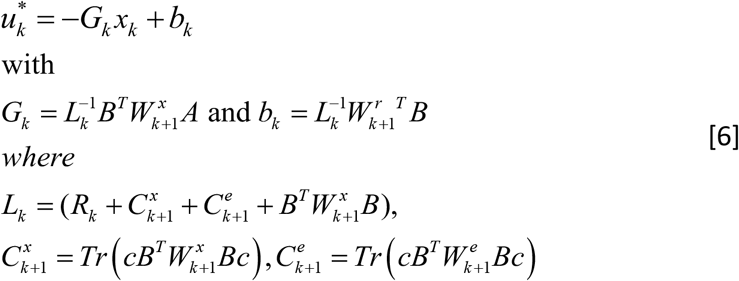

and the weights are calculated recursively backward till the first time-step using,

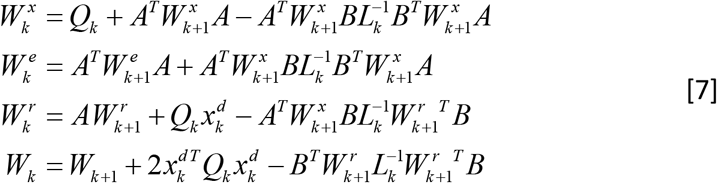

The weight matrices at the final step *n* are assumed to take the values 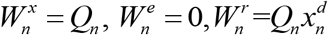 and 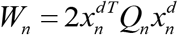. Since there is no true state feedback information available, the control signal in Eq. [6] is calculated on the estimated state 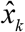 simultaneously propagating the forward model. The system dynamics which are given by Eq. [1] is used to calculate the mean trajectories of displacement and velocity. A detailed derivation of the model can be found in (Varsha et al. 2020, submitted manuscript).

#### Endpoint model used for comparison

The endpoint model is formulated as in (Shadmehr and Mussa-Ivaldi 2013), as a regulator problem. The main difference is that the cost function considered in this paper for the endpoint model is the same as in Eq. [3], with the same definition of the state *x_k_* as 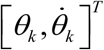 but the desired state 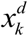 defined as 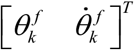, where 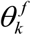 is the final saccade displacement to be achieved and 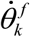 saccade displacement is taken as the mean saccade amplitude of every individual. Ideally, the final velocity is zero, but since we use a velocity threshold in the data processing, the end velocity has a non-zero value. This is accounted for by fixing 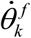 as the final value of the average velocity for every participant. The error weightage matrix *Q_k_* = [*q*_1_, 0; 0, *q*_2_] is modeled as a Heaviside function which is active only at the endpoint, where *q*_1_, *q*_2_ are free parameters. In this case, the optimal control signal 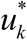 is obtained using the same procedure as used for the trajectory model but the value approximation used is 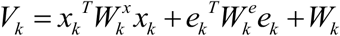. The optimal control is given by

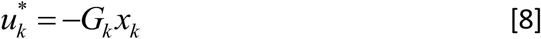

The *G_k_* definition is the same as in Eq. [6] while there is no bias term *b_k_*. The weight update equations are different and are given by the below expressions,

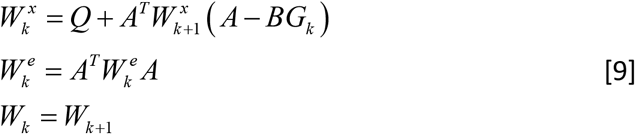

Note that the term with 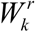 is absent in this case. The weight term *W_k_* remains constant since the time-step cost is not considered here unlike in (Shadmehr and Mussa-Ivaldi 2013). This assumption is used because the cost related to time-steps is suggested to be associated with the value assigned to the task and this remains the same across all trial types in our experiment. However, we have incorporated duration differences between the slow and fast trials into the formulation by simulating the model for the different durations observed in the experiment for slow and fast trials.

#### Model parameter estimation and validation

The velocity-based trajectory model has 2 free parameters namely *q* and *c*, while the final displacement-based endpoint model has 3 free parameters *q*_1_, *q*_2_ and *c*. To estimate the free parameters in the case of trajectory model, the mean data of angular velocity from the normal velocity block is used for each participant individually. The error weightage parameter *q* is estimated first with the noise parameter *c* set to zero. Further, with *q* fixed to the estimated value, the noise parameter is determined. In both cases, the normalized error as defined in Eq. [10] is minimized with *y* defined as angular velocity. The normalized error discussed above is defined for any variable of interest *y* as,

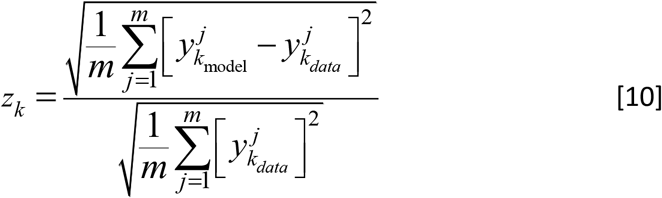

The initial guess value for this optimization was obtained from scanning of the parameter space based on values suggested in (Shadmehr and Mussa-Ivaldi 2013) for the endpoint model.

The same parameter estimation process was followed for the endpoint model. The endpoint model has three free parameters. The parameters *q*_1_ and *q*_2_ were estimated first, followed by *c*. However the normalized error function follows the same definition as in Eq. [10], but is calculated by considering *y* as the mean final displacement (i.e. the saccade amplitude). This version is referred to as the amplitude-tuned endpoint model. We also have another version of the endpoint model, which calculated the normalized error in Eq. [10] using the entire displacement profile. This version will be referred to as the displacement-tuned endpoint model. Note that all the figures in results present the amplitude-tuned endpoint model as this version best captures the philosophy of an endpoint model which has access to only desired amplitude information. However, the performance of the displacement-tuned endpoint model has also been discussed for each result.

The parameters obtained by this process from the normal velocity trials for each participant are used in the prediction of the mean angular displacement and mean angular velocity in their corresponding slow and fast velocity trials. Fit errors and prediction errors used for validation and comparison of models were quantified using the same formula as given in Eq. [10].

## RESULTS

### Model predictions of normative saccades

The velocity-based trajectory tracking stochastic optimal control model for saccade generation, which is referred to as the trajectory model, was evaluated for its ability to fit the experimental data. Data from the normal velocity trials was used. The desired velocity to be tracked by the system was assumed to be the mean velocity across trials for each participant. The fit obtained for mean angular velocity after estimating the two free parameters for an example participant is shown in Figure 3A.

**Figure 3.**
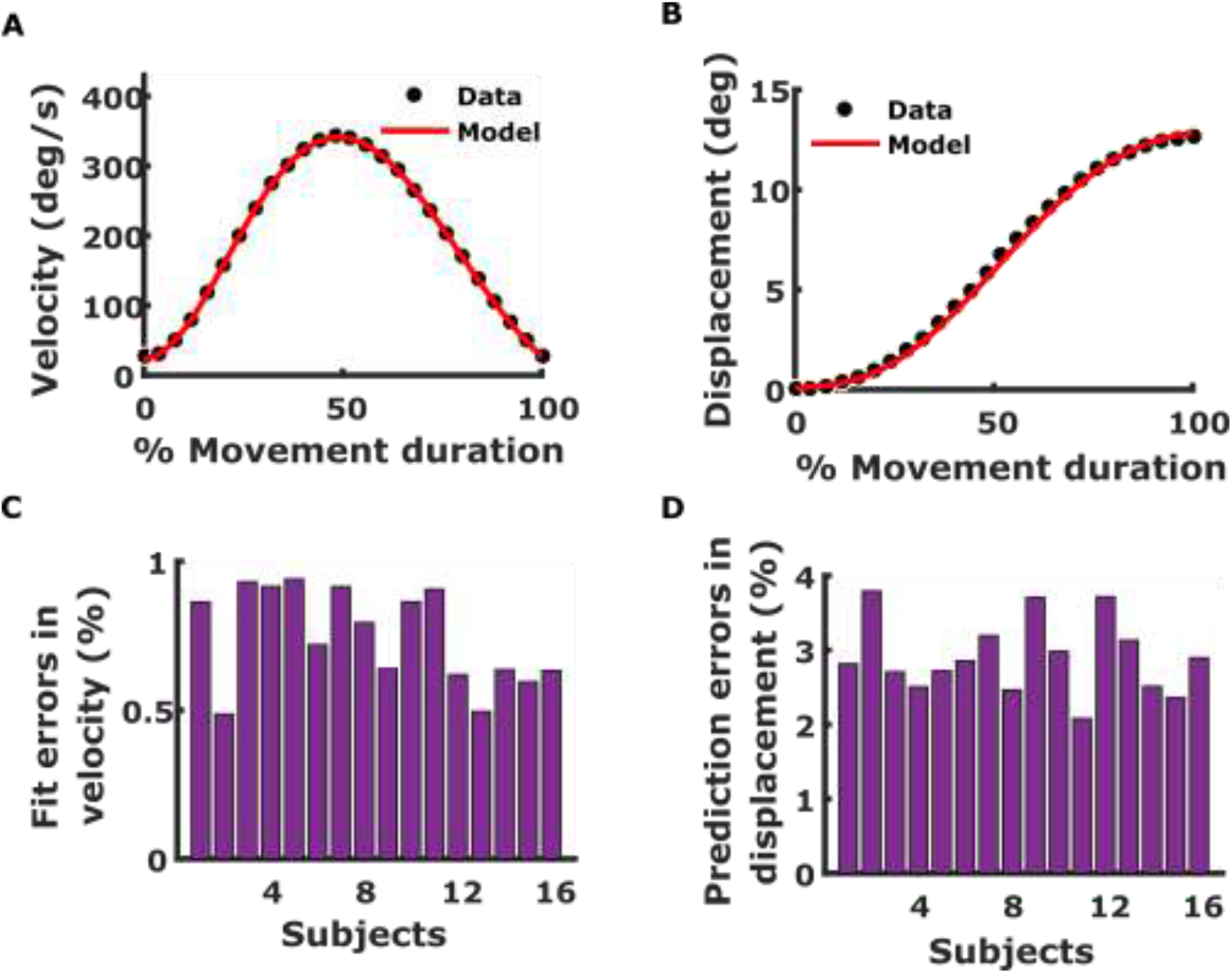
**A. Example fit of velocity data by the trajectory model.** The black dots markers represent the experimental data of angular velocity for a single subject and the red curve represents the simulation for the same by the trajectory model. **B. Example prediction of angular displacement by trajectory model.** Same as A but for angular displacement prediction. **C. Velocity fit errors across subjects.** Each bar gives the percent fit error for angular velocity for each subject. **D. Displacement prediction errors across subjects.** Same as C but for angular displacement prediction.

As expected, the velocity was well fit by the model. All fit errors in velocity were less than 1% with an average error across participants of 0.75 ± 0.16%. Crucially, although no information about the final displacement or displacement trajectory was given to the model, the displacement was also predicted well by the trajectory model. The average prediction error was 2.90 ± 0.50% across all participants.

The amplitude-tuned endpoint model’s (refer to model parameter estimation and validation section in methods) performance was also assessed. Its fit for the displacement data and its prediction of the angular velocity profile is shown in Figures 4A and 4B respectively for an example case.

**Figure 4.**
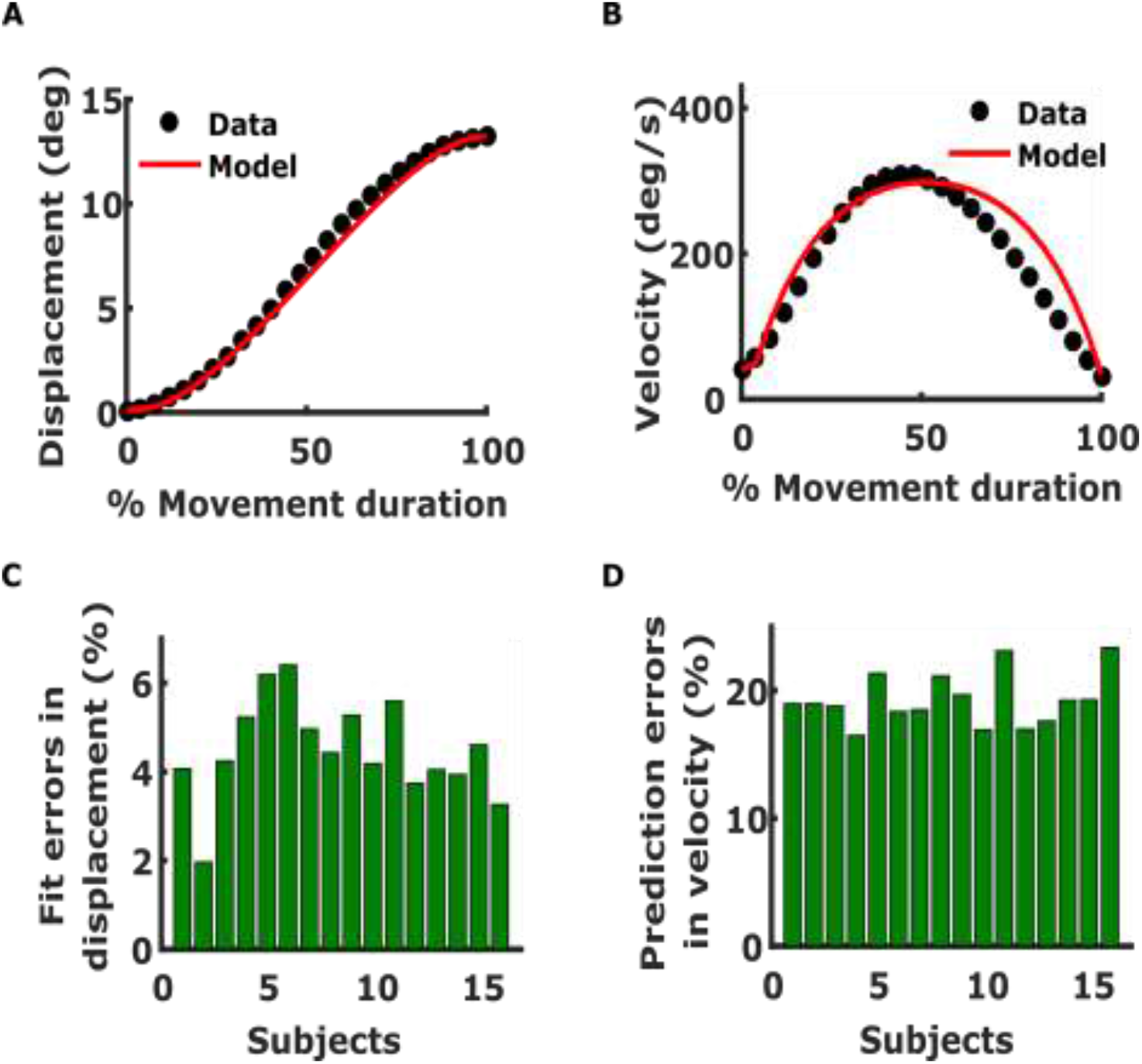
**A. Endpoint model example fit of displacement.** The black dot markers are the experimental data of angular displacement and the red curve is the simulation by the endpoint model. **B. Example prediction of angular velocity by the endpoint model.** Same as A but for angular velocity prediction. **C. Displacement fit errors across subjects.** Each bar gives the percent fit error for angular displacement for each subject. **D. Velocity prediction errors across subjects.** Same as C but for angular velocity prediction.

Across all subjects, the fit errors (4.51 ± 1.10%; Figure 4C) were observed to be higher than the prediction errors of the trajectory model for displacement (2.90 ± 0.50%; Figure 3D). However, the prediction of velocity by the amplitude-tuned endpoint model was bad with high errors of 19.34 ± 2.04% and was approximately twenty-five times the velocity fit errors of the trajectory model (0.75 ± 0.16%). Since the amplitude-tuned endpoint model did not fit the displacement well, we tested another version called the displacement-tuned endpoint model (refer to model parameter estimation and validation section in methods). In this version, the number of data-points used in parameter estimation was equal to the velocity-based trajectory model as it used the entire displacement profile. The displacement-tuned endpoint model produced better fits for the angular displacement (1.70 ± 1.30%) and had errors even lower than the displacement prediction errors of the trajectory model (Figure 3D). However, the displacement-tuned endpoint model still failed to improve the velocity fits. The predictions errors were high with an average of 16.24 ± 3.13% across the participants. This suggests that the trajectory model based on velocity is better in predicting the saccade velocity profiles compared to endpoint models.

### Modulation of saccade velocity in an eye-hand coordination task

The endpoint model suggests that the saccade velocity profile is generated by a controller that is solely based on the endpoint (amplitude) information. But there is evidence from prior research that suggests that saccadic velocity can be modulated independent of its amplitude (Gopal et al. 2017). This experimental evidence raises the possibility that the saccadic system engages an explicit velocity-based control system, as suggested by the trajectory model. To test this possibility, we reanalyzed the experimental data presented in (Gopal et al. 2017). We hypothesize that compared to the endpoint model; the velocity-based trajectory model would be able to fit better the observed behavior of saccades seen in the above experiment compared to the endpoint model. We assessed whether the trajectory model based on velocity could predict the behavior in the slow and fast velocity trials.

When the saccades associated with the slow and fast trials of hand movement were segregated, there was a significant difference in the mean peak velocity of these two subsets of saccade data. As also reported in (Gopal et al. 2017), to avoid the confound that the observed peak velocity differences may be due to differences in the mean amplitude of the saccades itself, we chose a subset of saccades that are amplitude matched for further analysis. In the example shown (Figure 5) we can see that even after amplitude matching (Figure 5A), there were significant differences in peak-velocities across slow and fast trials (Figure 5B). In comparison to the analysis done in (Gopal et al. 2017), we normalized the saccade duration (see methods) and found the results were consistent. We found that 10/16 participants showed significant differences in peak velocity (Figure 5D, p=0.0012, t (15) = 4) between fast (330.4 ± 39.8 degrees/s) and slow trials (311.6 ± 43.7 degrees/s) after amplitude matching (Figure 5C). We hypothesized that this amplitude-independent saccade velocity modulation would be predicted only by the trajectory model and not by the endpoint model.

**Figure 5.**
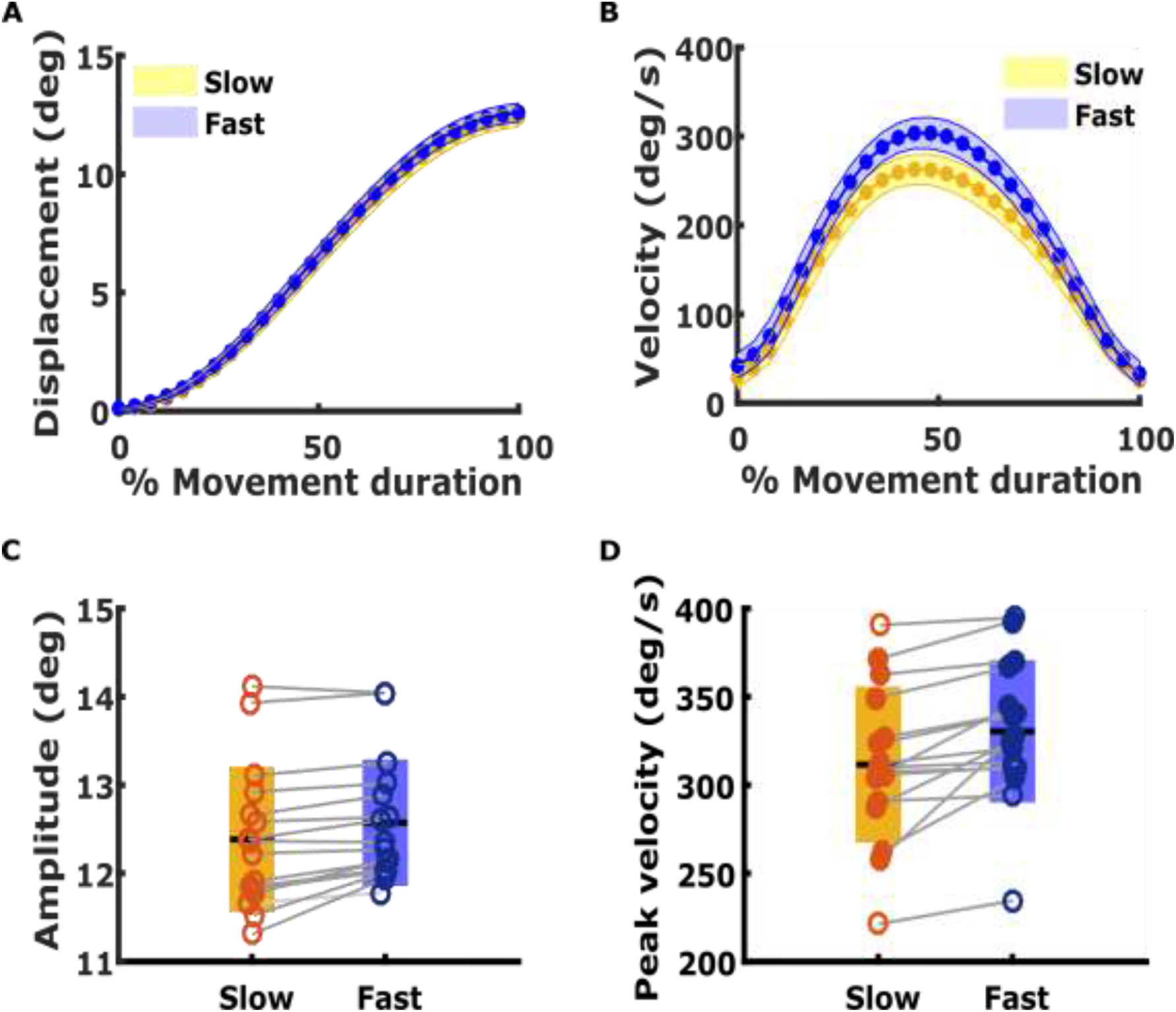
**A. Example displacement after amplitude matching and time normalisation.** Mean angular displacement of saccades in slow (yellow) and fast (blue) trials after amplitude matching are shown for an example subject. **B. Example velocity.** The mean velocity profiles for slow and fast trial of the same subject after amplitude matching and amplitude distribution are shown. Color coding is same as A **C. Comparison of saccade mean amplitudes.** The box plot shows the mean amplitudes of slow (yellow) and fast (blue) trials for all 16 subjects. Circular markers represent the mean value for individual subjects and unfilled circle means no significant difference between the trials. **D. Comparison of peak velocities.** Subjects showing significant peak velocity differences between slow and fast trials are represented with filled circles. All other conventions are same as in C.

### Model performance under velocity modulation

We tested the performances of the trajectory model and endpoint model in predicting saccade behavior in slow and fast trials. For both models, the parameters estimated from the normative saccades (normal velocity trials) were used in the prediction of displacement and velocity. However, the duration of simulation of saccades was fixed as the mean duration of all slow trials for the slow case and fast trials for the fast case. The prediction errors in displacement were significantly lower for the trajectory model compared to the amplitude-tuned endpoint model for slow trials (Figure 6A, p=0.001, t (15) = 4.06) as well as fast trials (Figure 6A, p=0.034, t (15) = 2.32). Contrary to this, the displacement-tuned endpoint model performed better than the trajectory model in predicting the displacement for both slow (p<0.001, t (15) = −5.33) and fast trials (p=0.009, t (15) = −2.95). However, in predicting the velocity of saccades the trajectory model performed better than the amplitude-tuned endpoint model in the slow trials (Figure 6B, p<0.001, W = 136) and fast trials (p<0.001, t (15) = 34), similar to the normative saccades. Even in the case of the displacement-tuned endpoint model, this trend remained the same with the trajectory model performing significantly better in predicting the velocity of slow (p,0.001, t (15) = 20.33) as well as fast trials (p<0.001, t (15) = 28.99).

**Figure 6.**
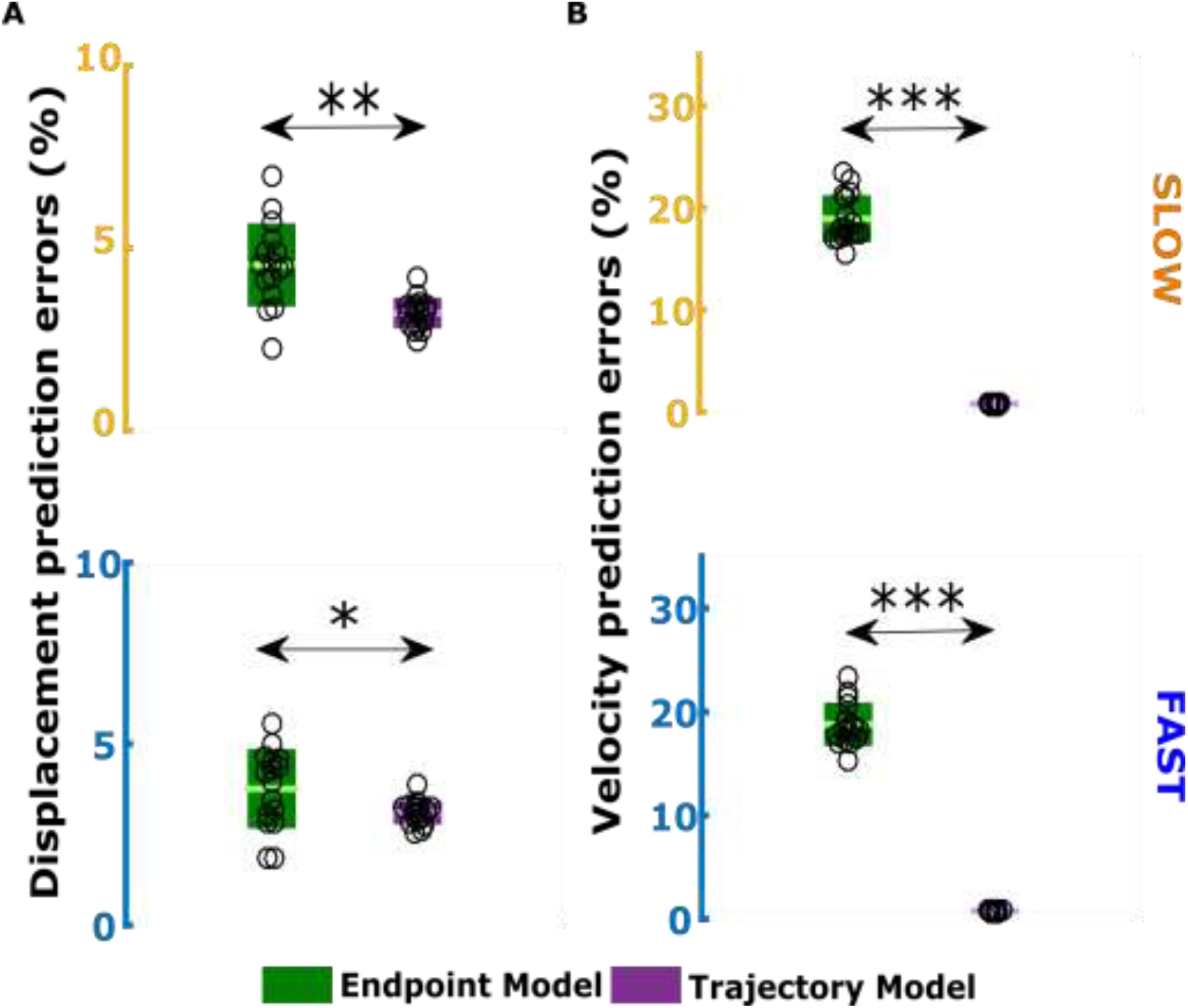
**A. Displacement prediction errors for slow (top) and fast (bottom) trials.** The plot shows a comparison of errors in prediction of displacement by the trajectory model (green) and endpoint model (violet). Each dot represents a subject and the central line on the bars indicate the mean value of fit errors across 16 subjects. The edges of the bars show one standard deviation above and below the mean. **B. Velocity fit errors for slow (top) and fast (bottom) trials.** The plot shows a comparison of prediction errors for the endpoint and trajectory models for velocity. The rest of the conventions are same as A.

Finally, we tested the endpoint model and the trajectory model for their ability to explain the amplitude matched peak velocity differences observed in the saccade behavior of slow and fast trials. The prediction of peak velocity differences by the trajectory model was closer to the experimental data than the endpoint model. The absolute errors in the prediction of peak velocity differences were calculated by taking the square root of the squared deviations between the experimental observation of peak velocity difference and model prediction of peak velocity difference for each participant. These absolute errors were quantified and compared (Figure. 7) for both the endpoint model (5.03 ± 5.51 degree/s) and trajectory model (0.04 ± 0.04 degree/s).

**Figure 7.**
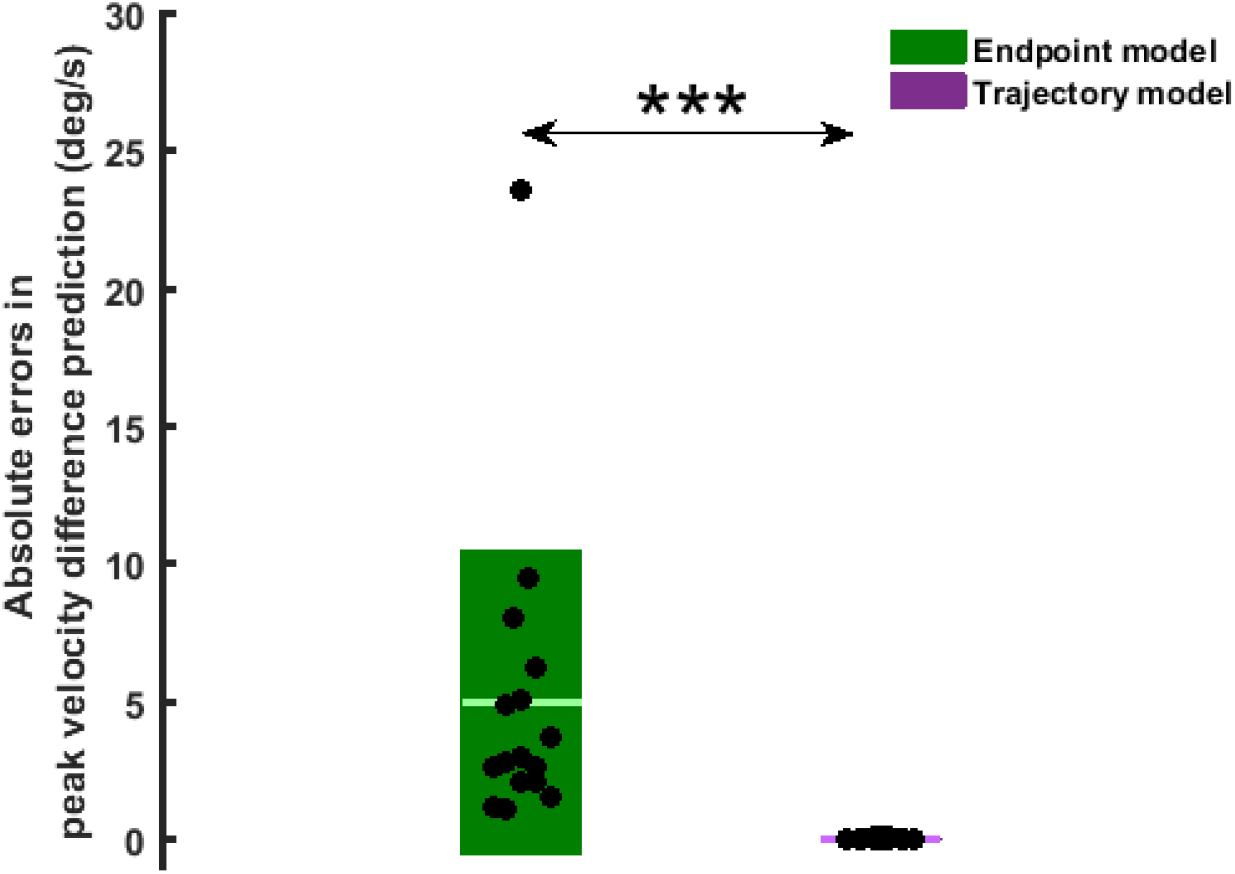
Peak velocity difference prediction errors by the models. The box plot shows the absolute errors in prediction of difference in the peak velocity between slow and fast trials for endpoint model (green) and trajectory model (violet) across subjects. The black dot represents errors for individual participants. The edges of the box represent one standard deviation from the mean errors represented by the central line. Note that the scales of errors of the velocity model prediction is so small compared to that endpoint model that the violet bar of the trajectory model is not visible.

The endpoint model failed to predict these peak velocity differences even though they could predict well the displacement profiles of the slow and fast trials. The predictions errors of the trajectory model based on velocity tracking were significantly smaller than the endpoint model (p<0.001, W = 136). The result remained unchanged even when the subject with large error for the endpoint model was left out during comparison. This result was also independent of whether the version of the endpoint model was amplitude-tuned or displacement-tuned. The endpoint model’s performance further degraded if the difference in duration between the slow and fast trials were not incorporated into the model simulations. Further, we also compared if the endpoint model and trajectory model performances were similar for the case of the 6 participants who did not show any significant difference in peak velocities between slow and fast trials after amplitude matching. It was observed that even in this subset of 6 participants, the velocity-based trajectory model’s predictions were significantly better than the endpoint model (p=0.009, t (15) = 4.13). These results suggest that only a trajectory control based on velocity can explain the saccadic behavior that is performed in conjunction with a hand movement, where there was an amplitude independent velocity modulation of saccades.

## DISCUSSION

Models of saccadic control have been predominantly explained by endpoint control, where the final displacement is the critical variable of interest. Such a control philosophy arose from the observations that input to the brainstem saccade generation system was encoded in the form of saccade amplitude and direction. In contrast, recent neurophysiological evidences propose the presence of a dynamic kinematic plan of velocity in the saccadic system, suggesting a possible trajectory planning of saccades (Goossens and van Opstal 2012; Smalianchuk et al. 2018). Here, we provide evidence of possible trajectory planning based control in the saccadic system that tracks the desired saccade velocity using a theoretical framework in which desired velocity is an explicitly tracked variable (Varsha et al. 2020, submitted manuscript). Not only did the velocity-based trajectory model fit normative saccade velocities better than the endpoint model, but this model also explained the saccadic velocities when it was modulated by the accompanying hand movements.

Most optimal control models are based on the philosophy that the saccade generation system is under endpoint control. Hence, these models try to attain the desired endpoint (amplitude) as accurately as possible (Harris and Wolpert 2006). If the trajectory of saccades is modulated depending on task goals or interruptions during the task, the saccadic system would need a controller that can keep a track of the desired trajectory rather than just the endpoint. The optimal feedback control model proposed for adaptive saccades indirectly suggested such an online control of saccades (Chen-Harris et al. 2008b). At each instant of time, the controller obtained a prediction of the states of the oculomotor plant from the forward model in the internal feedback loop. However, even in this optimal feedback model, the controller’s objective was to minimize the error between the final value of the states and the intermediate predicted states. The final value of states was taken as the saccade amplitude to be achieved and the end velocity (which was taken to be zero). Thus, there was no proposed explicit velocity plan incorporated in this endpoint model. This strategy seems valid, given the fact that saccades follow the main sequence relationship that interlinks the saccade amplitudes, duration, and peak velocity (Bahill et al. 1975). Thus, in contrast to having an explicit representation of velocity, it is plausible that the brain has a more abstract representation of velocity in terms of saccade duration and saccade amplitude. The tight coupling between the saccade amplitude and peak velocity suggests that the modulation in the saccade velocity trajectory can be achieved by planning for two different amplitudes of saccades. However, experimental evidence presented here suggests that the velocity modification can be independent of saccade amplitude; hence the control of saccades based on the endpoint amplitude plan alone is inadequate. This notion was reinforced by this study, which showed that the endpoint model underperformed relative to the trajectory model in predicting the difference in peak velocities between the amplitude matched slow and fast trials.

Can the differences in peak velocity observed experimentally arise because the saccades were planned to achieve the same amplitude, but with two different durations? Even this hypothesis is unlikely since, in the eye data analyzed, among the 10 participants who showed significant peak velocity differences between slow and fast trials, 7 had comparable duration distributions. Thus, at least in these participants, the peak velocity difference of saccades measured at the behavior level is probably because of the two different explicitly specified velocity plans. In a previous study, duration has been used as a variable in the cost function of the optimal feedback model based on the endpoint plan (Chen-Harris et al. 2008), in which the duration cost reflected the value associated with completing the task. However, in our experiment, the value of completing the task for both the slow and fast conditions of hand trials was the same, suggesting that duration need not be in the cost function. Despite this, we incorporated different durations into the model which improved the predictions of the endpoint model to some extent, but it still underperformed relative to the trajectory model in predicting mean velocities, and the peak velocity differences between amplitude matched slow and fast trials. Taken together, the endpoint model can handle variations within the normative main sequence constraint but fails to account for modulations in the velocity profile of saccades, independent of amplitude and duration changes.

Alternately, is it possible that to achieve the goal more accurately, the internal feedback loop of the brainstem saccade generating system causes the velocities to change between slow and fast trials instead of having two explicitly different saccade plans of velocity? Even this is unlikely since the average time at which the velocity profiles between the slow and fast trials diverged in our observed data was approximate ~20.5 ms; whereas the time required for internal feedback to kick in is estimated to be around 30 ms (West et al. 2009) when target information is available, 37 ms for predictive memory-guided saccades (Richardson et al. 2011) and greater than 45 ms for saccades when target information is not available during movement and during adaptive saccades (Xu-Wilson et al. 2011). This further strengthens the argument that the feedforward component of the saccade plan incorporates a representation of velocity.

The velocity-based trajectory model used in this work, for the saccadic system, incorporates an explicit plan in the form of the desired saccade velocity. The trajectory model was able to predict mean displacement and fit the mean velocity of normal saccades. At the same time, it out-performed the endpoint model in explaining amplitude independent velocity modulation in saccades in an eye-hand coordination task with velocity modulation instruction. This was observed even though the trajectory model based on velocity required only two free parameters compared to the endpoint model which had three free parameters. We also tried a version of the endpoint model (displacement-tuned) in which free parameters were assumed to be tuned based on the entire displacement profile rather than just the final displacement (amplitude-tuned). Neither the increased freedom in terms of free parameters nor the increased information used in parameter estimation could provide an advantage to the endpoint model in explaining saccade velocity data or its modulations. In general, based on its performance relative to the endpoint model, the velocity-based trajectory control is a feasible alternative.

The obvious question that arises is the neurophysiological basis of such an explicit velocity plan. Recently, (Smalianchuk et al. 2018) provided evidence for the presence of an explicit velocity control in the saccadic system by investigating the activity in the superior colliculus during normal and blink-perturbed saccades within single neurons. This is also congruent with previous work which showed that population activity of the cells in superior colliculus performs a dynamic ensemble coding that represents the intended movement trajectory (Goossens and van Opstal 2012; Goossens and Van Opstal 2006). Further, eye-hand coordination studies have observed modulation of saccadic velocity with hand movements (Van Donkelaar et al. 2004; Gopal et al. 2017; Snyder et al. 2002). Also, the activity in oculomotor structures like frontal eye fields and superior colliculus have been observed to change when eye movements are accompanied by hand movements (Reyes-Puerta et al. 2011; Thura et al. 2008). Based on these observations of kinematic interactions during eye-hand coordination, in conjunction with evidence that the neurons in the superior colliculus are sensitive to reach (Linzenbold and Himmelbach 2012; Stuphorn et al. 2017; Werner et al. 1997), we speculate that the neural instantiation of this velocity plan in case of eye-hand coordination also occurs in the superior colliculus.

Overall, data presented in this work as well as the trajectory model’s predictions, suggest that a saccade plan in the form of velocity encoding drives saccadic eye movements and their control. We suggest that this redundancy in the saccadic system to use a velocity-based control or endpoint control makes it robust and flexible to handle varying demands imposed by task context.

## AUTHOR CONTRIBUTIONS

VV- Varsha Vasudevan, AG- Atul Gopal, SJ- Sumitash Jana AM- Aditya Murthy, RP- Radhakant Padhi.

AG, SJ, and AM designed the experiments; AG and SJ conducted the experiments; VV and RP carried out the modeling of the data; VV, RP, and AM interpreted the results; VV prepared the figures and drafted the manuscript; SJ, AG, RP and AM edited and revised the final version of the manuscript.

